# Optimal marker genes for *c*-separated cell types

**DOI:** 10.1101/2025.02.12.637849

**Authors:** Bartol Borozan, Luka Borozan, Domagoj Ševerdija, Domagoj Matijević, Stefan Canzar

**Affiliations:** School of Applied Mathematics and Informatics, University of Osijek, Osijek, Croatia; Faculty of Informatics and Data Science, University of Regensburg, Regensburg, Germany

## Abstract

The identification of cell types in single-cell RNA-seq studies relies on the distinct expression signature of marker genes. A small set of target genes is also needed to design probes for targeted spatial transcriptomics experiments and to target proteins in single-cell spatial proteomics or for cell sorting. While traditional approaches have relied on testing one gene at a time for differential expression between a given cell type and the rest, more recent methods have highlighted the benefits of a joint selection of markers that together distinguish all pairs of cell types simultaneously. These combinatorial methods differ mostly in the notion of discrimination between cell types, using Euclidean or Manhattan distance in the dimensions of the selected marker genes, or the difference in the fraction of cells expressing a given marker gene. The resulting combinatorial optimization problems then seeks to identify a small set of genes that yield discrimination above a given threshold between all pairs of cell types. However, existing methods either consider all pairs of individual cells which becomes intractable even for medium-sized datasets, or ignore intra-cell type expression variation entirely by collapsing all cells of a given type to a single representative. Here we address these limitations and propose to find a small set of genes such that cell types are *c*-separated in the selected dimensions, a notion introduced previously in learning a mixture of Gaussians. To this end, we formulate a linear program that naturally takes into account expression variation within cell types without including each pair of individual cells in the model, leading to a highly stable set of marker genes that allow to accurately discriminate between cell types and that can be computed to optimality efficiently.

## 1 Introduction

A common first step in the analysis of single-cell data, such as those generated in atlases of the brain and other human organs and tissues [9, 2], is the characterization of the different cell types and their states based on the distinct expression of so-called marker genes. Typically, cells are first grouped according to their overall transcriptional similarity by a computational clustering method such as Seurat [6] and Scanpy [20]. Then genes are statistically tested for differential expression between one cluster and the rest (one-vs-rest) [6, 20], or between all pairs of clusters (package cran [12]). The detected marker genes are then used to label and biologically interpret cell clusters in an experimental context. Other general purpose statistical or machine learning based methods include the ANOVA F-test and decision trees [18]. Minimum-redundancy-maximum-relevance (mRMR) [16], on the other hand, does not only consider the relevance of a feature for an outcome but also takes into account redundancy among features. Similarly, ReliefF [10] uses nearest neighbors to indirectly account for feature dependencies. Other ranking based methods include SMaSH [13] and RankCorr [19] which have recently been benchmarked [17] against several other methods.

While a one-vs-rest comparison of gene expression allows to differentiate between major cell types, it might fail to detect genes whose expression differs only between similar subtypes, different tissues, or cell types in different disease states [7, 4]. In addition, such a cluster-by-cluster, gene-bygene approach will introduce redundancy into the set of selected genes, which makes it unsuitable for applications that rely on a small set of informative genes. For example, prominent fluorescence in situ hybridization (FISH)-based spatial transcriptomics technologies can assay only a limited number of genes that need to be pre-selected. Similarly, fluorescence-activated cell sorting (FACS) relies on a small number of distinct surface markers to sort cells by their types.

Rather than finding genes whose expression is distinct in a given cluster, recent methods have therefore studied a variant of the marker gene selection problem, which seeks to identify a single set of a small number of genes that jointly distinguish all clusters simultaneously. Existing methods model this task as a combinatorial optimization problem and differ mostly in how they define it or measure how much a given gene contributes to the discrimination between two clusters or cell types. scGeneFit [4] and G-PC [7] select marker genes such that distances between all pairs of cell types are above a certain threshold. The former method uses a linear program (LP) to minimize, for a fixed number of genes, the violation of the minimum required separation. The latter phrases the problem of minimizing the number of genes necessary to satisfy all distance requirements as a variant of the set cover problem which it solves using a greedy approach. Similarly, the method used in [11] uses set cover to minimize the number of genes to combinatorially distinguish a given cell type from all others in a single-nucleus RNA-seq data set of the mouse brain. It solves this set cover problem for each cell type using integer linear programming (ILP) and then uses a second ILP to integrate the resulting gene sets while reducing redundancy. To measure the separation between cell types, scGeneFit and G-PC use squared Euclidean or Manhattan distance, respectively, in the dimensions of the selected marker genes. Both methods compute the distance between single representatives (e.g. centroid) of each cell type. Alternatively, scGeneFit allows to explicitly model the separation of each pair of individual cells from two different types. The combinatorial method used in [11] does not take into account any form of distance between cell types but considers ‘distinguishable’ as a binary property, where each gene either distinguishes between two cell types or not, depending on the difference in the fraction of cells expressing it.

While one-vs-rest methods based on a statistical test of differential expression naturally take into account expression variation of a given gene across cells of the same type, combinatorial methods that aim to cover or separate all pairs of cell types with a single set of marker genes ignore within-cell type variation entirely by collapsing all cells of a given type to a single representative. On the other hand, including all pairs of individual cells in an LP, as scGeneFit does, becomes intractable even for small- and medium-sized datasets. It therefore selects a small set of constraints (5000 by default) among typically many millions of possible cell pairs at random, thus ignoring large parts of the generated data. Here we address these limitations and propose a combinatorial method that takes into account gene expression variation. Our method’s objective is to find a small set of genes such that cell types are *c*-separated in the selected dimensions. In previous work [3], two Gaussians have been defined to be *c*-separated if their centers are *c* radii apart. Random projection was then used to reduce dimensionality while approximately retaining the separation of a mixture of Gaussians. Instead, here we formulate an ILP based on a linear approximation that systematically searches for such a subspace without making the assumption of normality. We provide experimental evidence that this strategy yields marker genes that more accurately discriminate between distinct and similar cell types.

## 2 Methods

Our approach exploits the notion introduced in [3] that formalizes that a mixture of Gaussians is *c*-separated if its component Gaussians are pairwise *c*-separated.

### Definition 1.

Let *N* (*µ*_1_, Σ_1_) and *N* (*µ*_2_, Σ_2_) represent two Gaussians in ℝ^*d*^, where *µ*_1_ and *µ*_2_ are the means, and Σ_1_, Σ_2_ are the covariance matrices, and let tr() denote the trace of a matrix. We say that two Gaussians are *c*-separated if

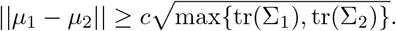

The motivation in [3] for choosing 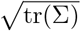 as the target separation of the means in the above definition is that tr(Σ) is the expected squared Euclidean distance from the mean:

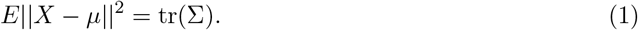

Furthermore, for a Gaussian with bounded eccentricity, where eccentricity was used in [3] to measure how non-spherical a Gaussian is, the distribution will be concentrated around this *radius* of 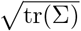. The degree of separation is controlled by parameter *c*. Larger values of *c* imply stricter separation with less overlap. A 2-separated mixture will be almost entirely separated, while 1- or 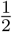-separated clusters will have a small overlap [3]. Single-cell count data restricted to a single cell type typically do not follow a Gaussian distribution [15]. However, as noted in [3], random projections of many high-dimensional distributions look more Gaussian due to the central limit theorem. In fact, in the marker gene subspace we observe (Figure 1) that most of the probability mass lies close to distance 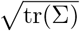 from the mean, rendering Definition 1 suitable as a target separation between cell types.

**Figure 1:**
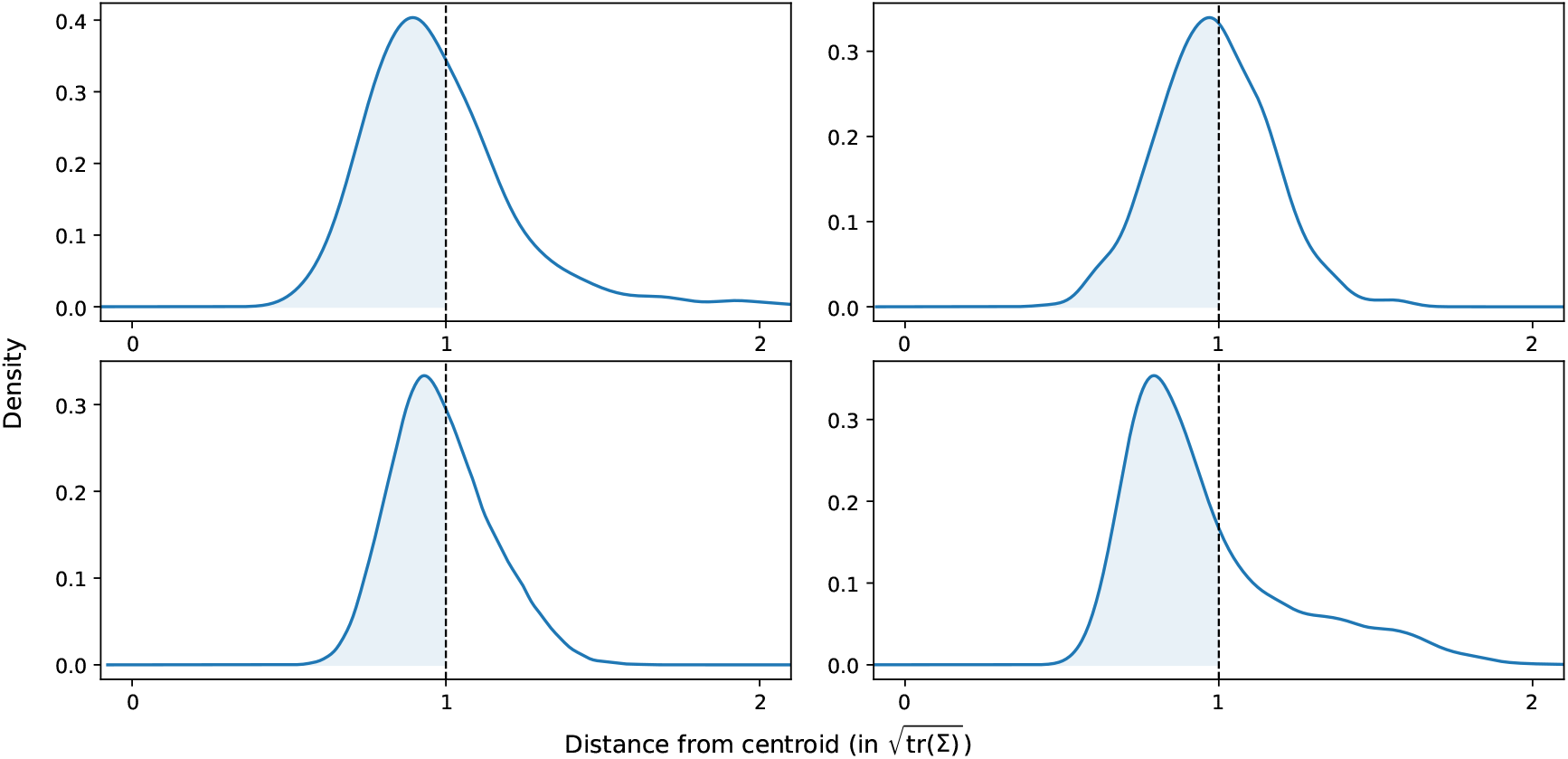
Probability mass of the distances to the centers (sample means) in units of 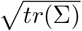 for the two largest cell types in the Zheng8eq (top) and IPF (bottom) datasets. The fraction of data lying within one standard deviation of the mean is reported as 63%, 55%, 57%, and 68%, from left to right and top to bottom.

### 2.1 c-separated point sets in *α*-subspace

Let *S* denote a sample of points in ℝ^*d*^. With a slight abuse of notation, below we do not distinguish between population parameters and (estimated) sample statistics. Analogous to Definition 1 we can compute the total sample variance tr(Σ) of *S* in the following way:

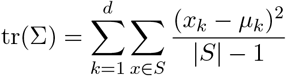

where *µ* denotes the sample mean of set *S, k* stands for the *k*-th component of a vector in ℝ^*d*^ and |*S*| denotes the cardinality of *S*. Let *S*^*α*^ denote the set of points induced by a vector *α* ∈ {0, 1}^*d*^, such that

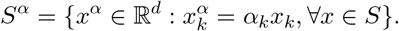

Namely, *α*_*k*_ = 1 denotes that the *k*-th coordinate is included, while *α*_*k*_ = 0 denotes that the *k*-th coordinate is ignored. We consider all coordinates for which *α*_*k*_ = 1 to define a point in a *α*-subspace. Then, the trace of the sample covariance in the *α*-subspace can be computed as follows:

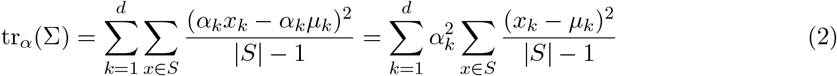

Now, for two samples of points *S*_1_ and *S*_2_, we are ready to rewrite the inequality from Definition 1 in some arbitrary *α*-subspace:

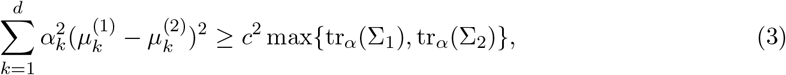

where *µ*^(1)^, *µ*^(2)^ denote the sample means of *S*_1_ and *S*_2_, respectively, and tr_*α*_(Σ_1_) and tr_*α*_(Σ_2_) are computed as in (2), for sample covariance matrices Σ_1_, Σ_2_ of *S*_1_ and *S*_2_, respectively. We call any two point sets *S*_1_ and *S*_2_ in ℝ^*d*^ *c*-separated in *α*-subspace if (3) holds.

### 2.2 c-separation as optimization problem

To identify an *α*-subspace of fixed dimension *m* such that *m < d*, where cells of different types achieve *c*-separation, we formulate a corresponding constrained optimization problem. Specifically, the goal is to minimize a linear objective function that imposes a penalty when the separation of cell types within the *α*-subspace is not fully realized, subject to a set of nonlinear constraints ensuring distinct separation properties for each cell type. To enable tractable computation, we further linearize these constraints locally around a chosen point, approximating the original problem as an instance of an Integer Linear Program (ILP).

Given input point set *S* ⊂ ℝ^*d*^, let *S*_*i*_ ⊆ *S, i* = 1, …, *l* denote all cells of label *i*. For every pair of sets *S*_*i*_ and *S*_*j*_, we aim to separate them according to (3). For an arbitrary value of *c* ∈ ℝ, finding an *α*-subspace such that all constraints (3) are satisfied will in general not be feasible. Therefore, we rephrase (3) by introducing slack variables *β* as follows:

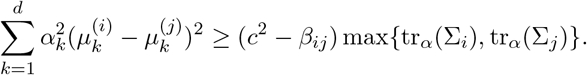

By increasing values of the slack variables *β*_*ij*_, we allow for a relaxation of the original requirement that the two sets *S*_*i*_ and *S*_*j*_ must be *c*-separated. Let *L* = {{*i, j*} : *i* = 1, …, *l, j* = 1, …, *l, i ≠ j*} be the set of all unordered pairs of cell labels. We can now formulate the search for an appropriate *α*-subspace as the following optimization problem:

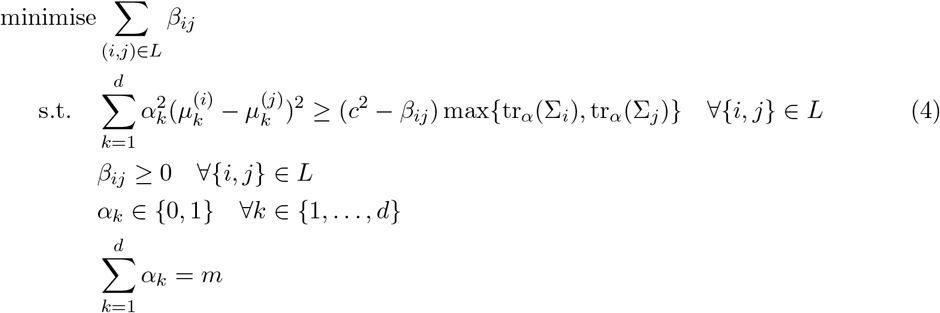

Note that *m* is the dimension of the subspace induced by the vector *α*. Furthermore, since *α*_*k*_ can only take the values zero or one, we can substitute 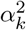 with *α*_*k*_ in (4). Constraints (4) are not linear since variables *β*_*ij*_ are multiplied with the trace of the covariance matrix of either *S*_*i*_ or *S*_*j*_ that in turn depend on variables *α*. We will linearize them in the next Section.

#### 2.2.1 Integer Linear Program (ILP) formulation

Let 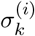, the *k*-th element on the diagonal of the matrix 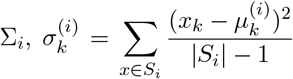, denote the sample variance for gene *k* in *S*_*i*_. For a fixed *α* ∈ {0, 1}^*d*^, the trace of the covariance matrix Σ_*i*_ can be calculated as 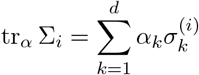. Thus, constraint (4) can be written as:

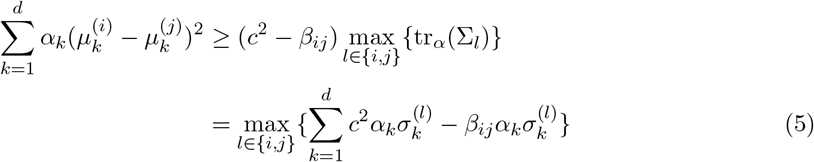

To linearize equation (5), we need to eliminate the product of the variables *β*_*ij*_ and *α*_*k*_, as both are part of our optimization problem. Given that *β*_*ij*_ ∈ [0, *c*^2^] and *α*_*k*_ ∈ [0, 1], we can linearize their product around the midpoint 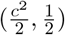 by employing the first-order Taylor expansion of the function *f* (*β*_*ij*_, *α*_*k*_) = *β*_*ij*_*α*_*k*_ at the point 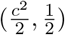:

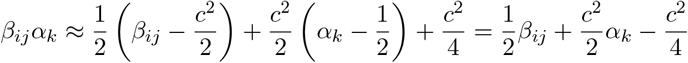

When we linearize around the midpoint, the linear approximation may deviate from the true product value at the boundaries of the intervals. The worst-case scenario arises when both variables reach their extreme values, which leads to an upper bound on the error of 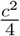. Conversely, the expected deviation can be calculated as the average of the errors over the range of possible values. Since the linearization occurs around the midpoint, the average error is 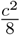. All our experiments will be conducted with *c* = 0.4 (see Section 3), which yields small errors in practice. Continuing (5), we get:

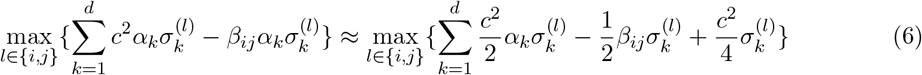

The above optimization problem can now be written in linear form as:

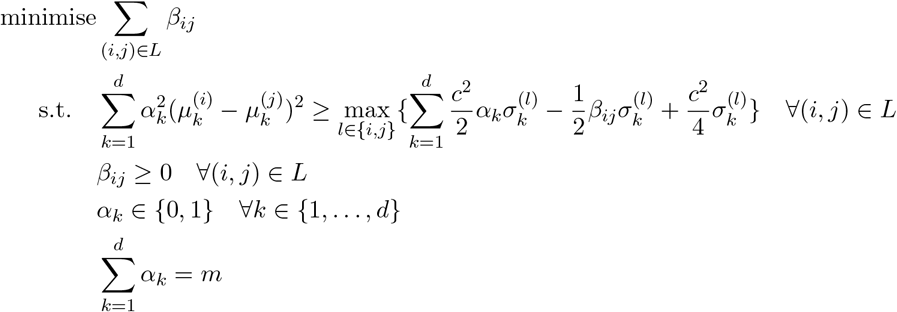

By introducing new slack variables *z*_*ij*_ to eliminate the maximum function, we can finally express the previous problem as an integer linear program (ILP):

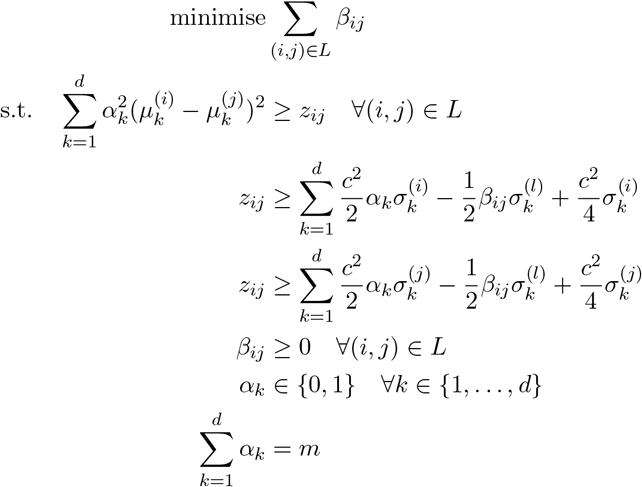

## 3 Results

We implemented the ILP described in the previous Section in method SepSolve. We used Gurobi to solve its linear relaxation and convert fractional *α* to integral values through a simple ranking method which rounds the *m* largest values to 1 and sets the remaining *α*-variables to zero. The same rounding scheme was applied in [4]. We compared the performance of SepSolve to most recent combinatorial methods scGeneFit [4] and G-PC [7]. No implementation of the combinatorial method used in [11] was available. In [7], G-PC was compared to eight previous methods and, along with scGeneFit, demonstrated the best performance on a subset of the datasets used in this study. Most of the other methods mentioned in the Introduction were recently benchmarked in [17], with combinatorial methods excluded as out of scope. All experiments were run on an AMD Ryzen Threadripper 3990X 64-Core Processor @ 4.3GHz, 256 GB of RAM, and Python version 3.10.12.

All methods, including SepSolve, were run using default values for all parameters. In SepSolve we set the default value for the separation constant *c* to 0.4. On smaller datasets, scGeneFit was run in both pairwise and centers mode, including constraints for all pairs of cells or only between single representatives (centers) for each cell type, respectively. The authors’ experiments have shown that the centers mode is the most efficient and stable one. Indeed, attempts to run scGeneFit in pairwise mode on the larger datasets failed because it exceeded available memory or a time limit of 12 hours. The authors of G-PC did not provide a default value for the coverage factor (*K*) in their implementation. Consistent with [7], we therefore varied *K* in the range of {5, …, 20} and chose the value that yielded the highest average accuracy on the IPF dataset. The same value was then applied to all datasets.

### 3.1 Proof of principle

To illustrate the benefits of taking variance into account in the combinatorial model, we have created a synthetic three-dimensional spherical dataset consisting of four different labels (i.e. cell types) and three genes A, B and C. The mean expression levels of gene A do not differ much between the first two cell types and between the last two, with a difference of only 0.45 units. However, across all cell types, the variance of expression levels of gene A is low, namely 0.50. The expression means of genes B and C are equidistantly distributed, with a relatively large distance of 8 between neighboring cell types. However, the expression of genes B and C exhibit a high variance of 12 across all cell types. As illustrated in Fig. 2, the marker gene selection methods scGeneFit and G-PC, which ignore gene expression variation, choose genes B and C. SepSolve, on the other hand, leverages very small variation in the expression of gene A and selects it as a marker, yielding a much sharper separation of cell types.

**Figure 2:**
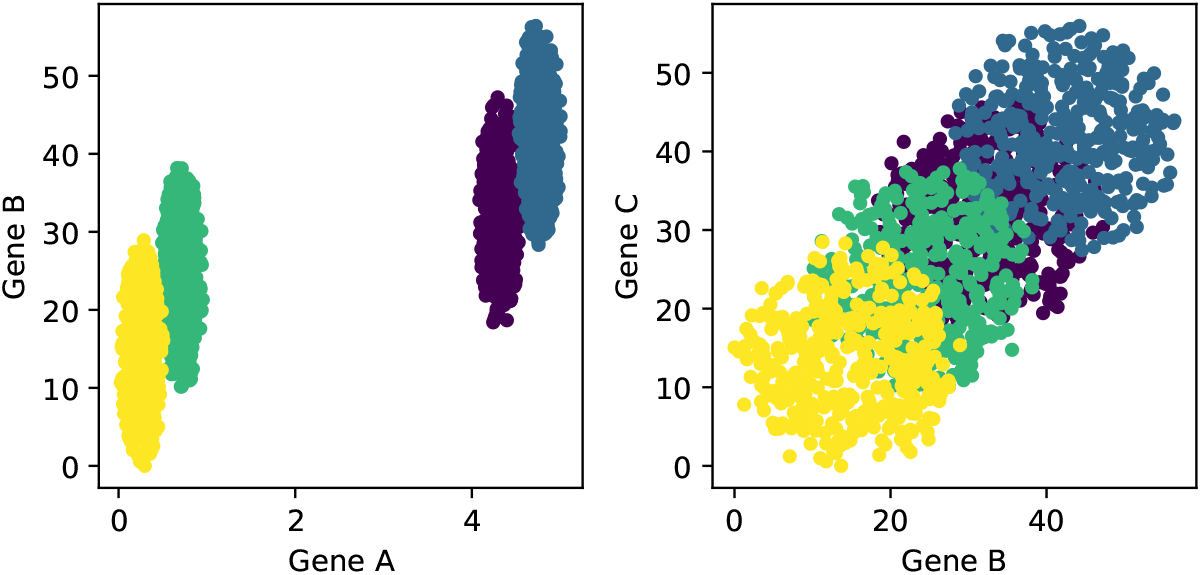
Cells from four different types in the projected space of marker genes computed by SepSolver (left) and scGeneFit and G-PC (right).

### 3.2 Classification accuracy

Similar to the benchmarks performed in [4] and [7], we evaluated the discriminatory properties of the identified marker sets by the ability of a classifier to predict cell types based on them. We benchmarked methods on four different scRNA-seq datasets (Table 1). We included all three datasets used in [7] from the lung, mouse cortex, and a human cell atlas. Consistent with [7], we restricted the IPF dataset to the healthy samples. We used an additional dataset (zheng8eq) of peripheral blood mononuclear cells (PBMCs) in our benchmark which included both distinct and similar cell types [14]. True cell-type labels were taken from the original publications. In the zheng8eq dataset the authors randomly mixed roughly equal proportions of presorted B cells, CD14 monocytes, naive cytotoxic T cells, regulatory T cells, CD56 NK cells, memory T cells, CD4 T helper cells, and naive T cells. In this case, true labels were independent of scRNA-seq measurements.

**Table 1:** Datasets used in this study.

| Dataset | #Cells | #Genes | #Cell Types | Reference |
| --- | --- | --- | --- | --- |
| IPF (Idiopathic pulmonary fibrosis) | 96,303 | 45,947 | 38 | Adams et al. [1] |
| HCA (Human Cell Atlas) | 84,363 | 22,653 | 24 | He et al. [8] |
| Mouse cortex | 3,005 | 20,006 | 7 | Zeisel et al. [21] |
| Zheng8eq | 3,971 | 1,571 | 8 | Duò et al. [5] |

Following the same preprocessing steps as applied in [7], we removed cell types with less than 50 cells and genes expressed in less than 10 cells. Counts were normalized to 10,000 using the scanpy python package and log transformed after adding a pseudocount of 1. Finally, we identified highly variable genes using the highly variable genes function from Scanpy, applying the same parameters as those used in [7] for all datasets. Following the authors’ recommendation, we have additionally standardized the data to zero mean and unit variance when running G-PC.

Consistent with the analysis in [4] and [7], we used the marker genes found by each method in k-nearest neighbors and logistic regression classifiers. For the logistic regression classifier we split the data into 70% for training and 30% for testing. As in [7], we measured the performance of the classifiers using the macro-average F1 score which corrects for an unbalanced cell type composition and gives higher weight to rare cell types. It is computed as the mean F1 score across all cell types.

Fig. 3 demonstrates an improved ability of the logistic regression classifier to predict cell type labels based on marker genes found by SepSolve. The increase in F1 score is particularly pronounced for a smaller number of marker genes on the HCA and the mouse cortex data, were F1 scores converge to near identical values for larger numbers of markers. On the zheng8eq dataset, a substantial improvement is achieved by SepSolve across the entire range of marker genes. The smallest improvement can be observed on the IPF dataset. However, it is the only dataset were a slight improvement can be observed even for larger numbers of markers. When selecting 160 marker genes, for example, SepSolve’s gene set yielded a F1 score of 86%, compared to a score of 83% and 84% obtained based on scGeneFit’s and G-PC’s markers, respectively. Using a k-NN classifier instead of logistic regression yielded similar results (Appendix, Figure S1).

**Figure 3:**
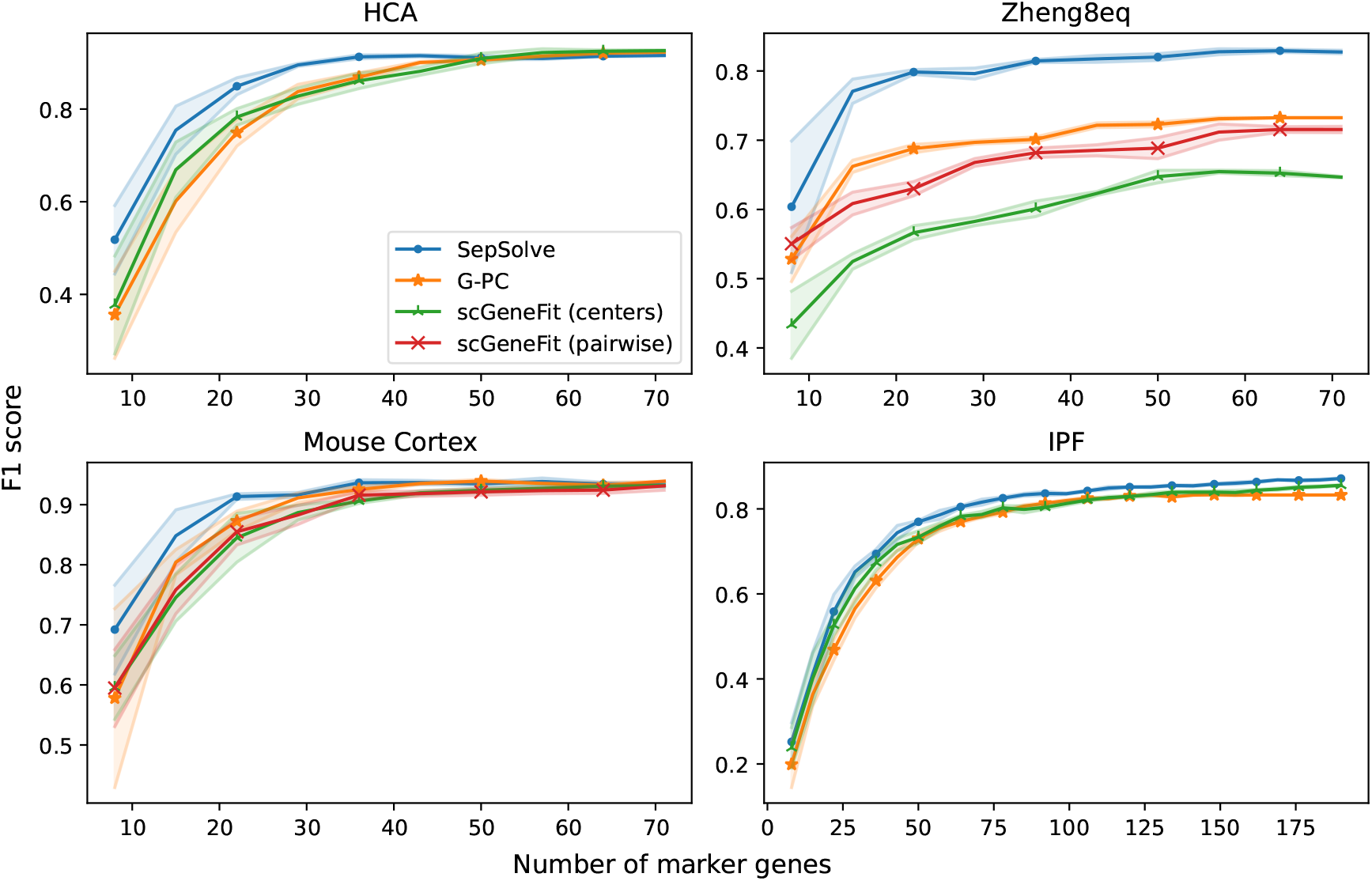
F1 scores of a logistic regression classifier when provided varying numbers of marker genes computed by the different methods on the four datasets. scGeneFit was run in pairwise and centers mode (see text for details). Shaded regions depict standard deviation.

### 3.3 Stability

In the context of feature selection, stability refers to the degree of consistency with which a feature is identified as relevant across different conditions. To be practical–such as when designing probes for a targeted spatial transcriptomics experiment based on a reference scRNA-seq sample–a set of marker genes must not only accurately discriminate between cell types but also remain stable across varying experimental conditions, including the number of profiled cells and different noise levels. We follow the approach in [7] and calculate the stability of a collection of marker gene sets {*S*_1_, *S*_2_, …, *S*_*n*_} as the average Jaccard similarity of all pairs of sets:

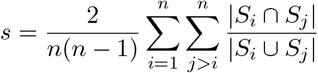

We evaluated the stability of markers computed by the different methods in two experiments. To avoid a potential confounding influence of the robustness of highly variable gene (hvg) detection on overall stability, we omitted the selection of hvgs in these experiments. First, consistent with [7], we randomly sampled 50% of input cells in 5 random trials. We ran scGeneFit, G-PC, and SepSolve on each subsample and then calculated the stability of the marker gene sets found by each method. Fig. 4 (left) shows that SepSolve returns a highly stable set of marker genes with around 80% overlap across all four datasets. The slightly higher stability observed on datasets IPF and HCA might be due to the large number of profiled cells. In contrast, both scGeneFit and G-PC achieved comparable stability only on a single dataset, mouse cortex in the case of scGeneFit and zheng8eq in the case of G-PC. In the worst case, only half or less than half of marker genes returned by scGeneFit and G-PC overlapped across runs. In addition, we assessed stability with respect to a small data perturbation. To this end, for each entry *e* in the raw count matrix, we sampled from a Poisson distribution with expectation 0.05 · *e* and either added or subtracted the obtained value from the matrix entry with equal probability. Again, this perturbation was applied 5 times, and the stability of detected markers was evaluated. Marker genes found by SepSolve were not much affected by this perturbation and overlapped by more than 95% across datasets. Marker gene sets returned by scGeneFit and G-PC where less stable across all datasets but were overall less affected than in the subsampling experiment. With 64% and 75%, they achieved the lowest stability values on the IPF dataset.

**Figure 4:**
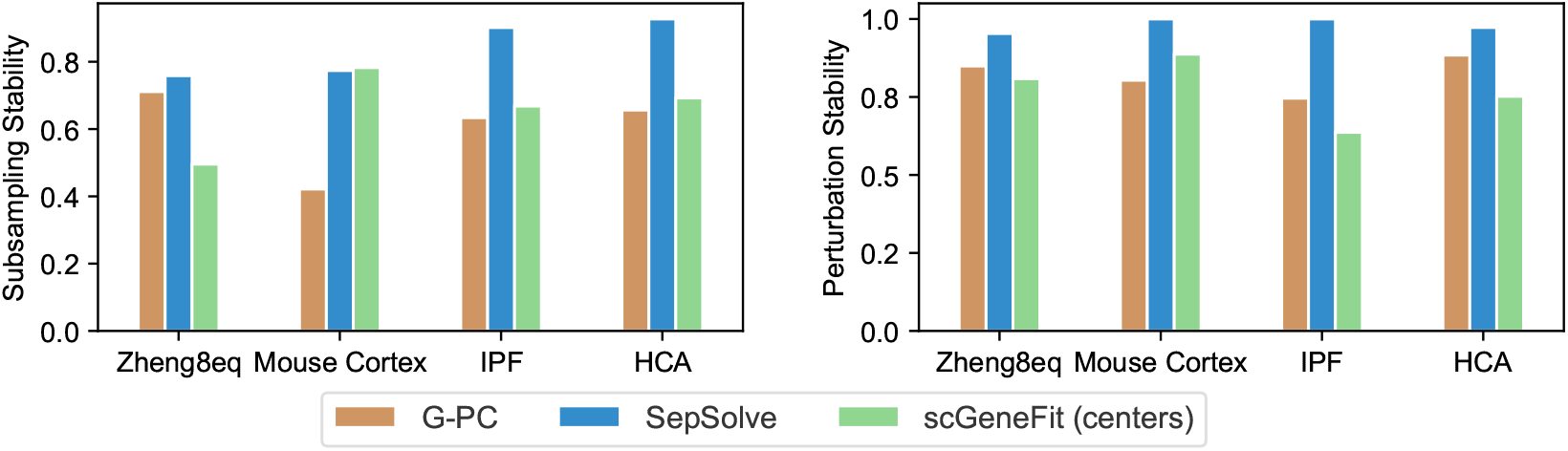
Stability of 50 marker genes computed by the different methods on random subsamples of cells (left) or perturbed counts (right).

### 3.4 Running times

Figure 5 displays the average running times of the different methods across all numbers of target marker genes evaluated in Figure 3 for the four datasets. Running times exhibit minimal variability with respect to the number of target genes (data not shown). As expected, G-PC’s greedy heuristic is the fastest and takes at most 1.23 seconds on the largest dataset (IPF). Solving a linear program and subsequent rounding in SepSolve took only 7.86 seconds in the worst case, while scGeneFit, run in the more efficient centers mode, took up to 52.93 seconds to complete. In pairwise mode, the running time of scGeneFit was 82.6 seconds on the smaller mouse cortex data. In this mode, scGeneFit failed to run on the larger datasets due to memory and time limits (12h).

**Figure 5:**
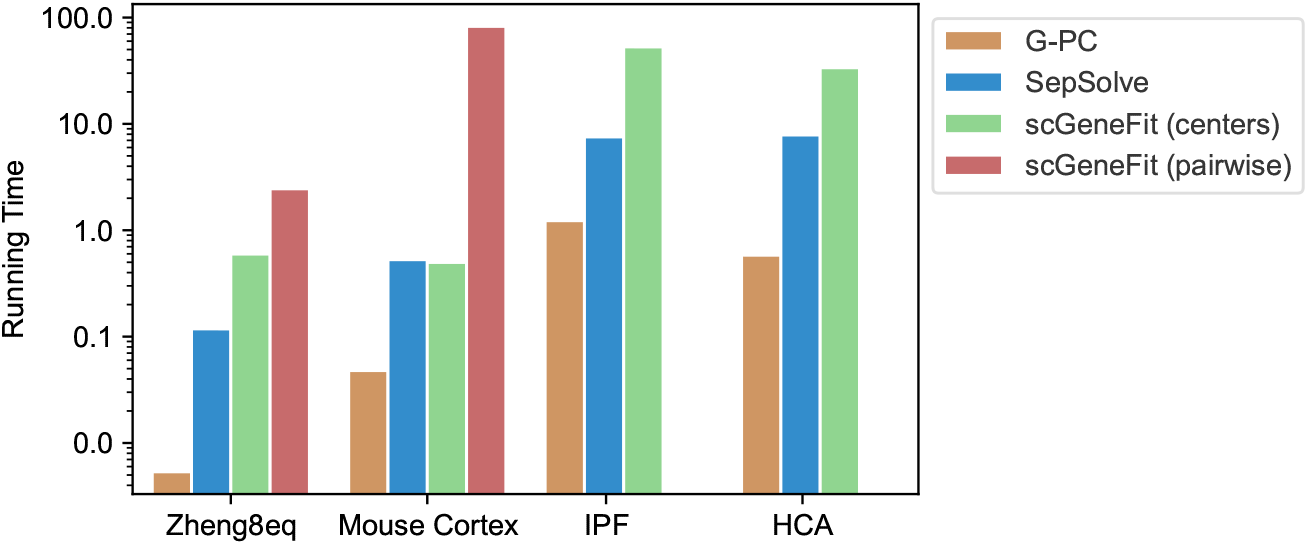
Average running times (log scale, in seconds) over all evaluated target numbers of marker genes on the four datasets.

## 4 Conclusion

We have proposed method SepSolve, a combinatorial approach that tries to discriminate between all cell types simultaneously in the space of marker genes. In contrast to existing methods in this class, it takes into account gene expression variability. Transcriptional heterogeneity within cell types can be due to different, potentially unknown, cell states and transitions between them, but can also have technical reasons such as noisy experimental measurements.

SepSolve annotated cell types more accurately, especially for a small number of marker genes which is the most common application scenario, for example when designing probes for a targeted spatial transcriptomics assay. Notably, scGeneFit achieved substantial improvements in the F1 score on the zheng8eq dataset when including constraints for all pairs of cells instead of single representatives only, into an otherwise identical optimization model. This again indicates the value of the additional information contained in the within-cell-type expression variation that SepSolve leverages effectively.

An additional advantage of our method is that it provides the most stable set of markers. This is a particularly useful property, for example, when designing spatial probes based on dissociated reference data. Again, we attribute this mainly to taking into account transcriptional heterogeneity within cell types in our model without including every pair of individual cells as scGeneFit does in the pairwise mode. In addition, we optimally solve an LP formulation of the problem, compared to the greedy scheme applied by G-PC that might be more sensitive to small data perturbations [7]. Furthermore, even though SepSolve takes into account within-cell-type variability, it is fast and can be applied to very large datasets. This is in contrast to scGeneFit, especially when trying to account for expression variability in pairwise mode.

Our method belongs to the class of filter methods that do not optimize a specific classifier during feature selection, in contrast to wrapper and embedded methods [7]. The low-dimensional subspace in which we represent cell cluster structure can therefore be useful for other tasks such as the deconvolution of bulk expression data as illustrated in [7].

A current limitation of our method is that total expression variance, i.e. the trace of the sample covariance, is used as target separation between cell types, inspired by previous work [3] on random projections of high-dimensional Gaussians. Even though we empirically showed that this target separation is justified for single-cell data, more tailored measures might exist. The flexibility of an LP formulation will allow us to investigate alternative measures of separation in future work. A second limitation is the simple ranking scheme we applied to convert fractional solutions of the LP to an integral one. We will explore more accurate approaches such as branch and bound with heuristic speed-ups and (randomized) rounding schemes with potential guarantees in the future.

SepSolve is available at https://github.com/bborozan/SepSolve.

## Notes

### Competing Interest Statement

The authors have declared no competing interest.

